# Germline-limited transposon remnants foster somatic genome diversification

**DOI:** 10.1101/2025.11.02.686105

**Authors:** Bozhidar-Adrian Stefanov, Mariusz Nowacki

## Abstract

Transposable elements are mobile DNA sequences that propagate within genomes, often with deleterious effects on host fitness. Some organisms excise transposons from their somatic genomes via programmed DNA elimination yet paradoxically retain them in their germline. The evolutionary rationale for this selective retention remains poorly understood. To address this, we first set out to develop a germline enrichment method in *Paramecium tetraurelia*, since the somatic DNA is several hundred times more abundant than the transposon-rich germline. Our approach improved the germline assembly significantly by reducing the number of contigs from the previously published 26186 to just 179, close to the expected chromosomal count. Contrary to the view that the germline serves merely for transgenerational genome propagation, long-read sequencing of poly(A)+ RNA uncovered abundant coding transcripts derived from germline-limited DNA. Furthermore, we detected numerous modified somatic transcripts caused by incorporation of ultrashort transposon remnants from germline sequences. Notably, these insertions show a size bias that preserves open reading frames, consistent with translation mediated quality control. Together, our results suggest that the germ-soma nuclear dimorphism serves as an intrinsic system for the emergence of genome innovations and transcriptome diversification through the retention of germline-limited sequences.

## Introduction

The evolution of genomes represents a dynamic process driven by the accumulation of deleterious, neutral, and advantageous mutations. A substantial fraction of these genetic alterations arises from faulty repair following DNA damage, or by single-base substitutions during DNA replication. Interestingly, transcriptionally active sites accumulate more base substitutions, presumably due to increased chromatin accessibility^1^ or the molecular forces excreted by RNA and DNA polymerases^2,3^. Since purifying selection^4^ efficiently reduces deleterious mutations, neutral variants constitute the predominant source of heritable genetic variation^5^. This fosters adaptability and improves responsiveness to changing environments and can confer a selective advantage over a static, non-evolving genome^6,7^. Base-substitution rates vary across lineages^8,9^, likely reflecting differing evolutionary pressure for adaptation and the effective population sizes^10,11^. Among the lowest mutation rates observed in eukaryotes was found in the protozoan species *Paramecium tetraurelia*^12,13^. This can be attributed to specialized polymerases^12,14^ combined with a nuclear dimorphism, in which a dedicated germline nucleus safeguards genetic information across generations while the gene expression is tasked to a separate somatic nucleus^15^. A low substitution rate does not preclude genomic evolution, since other mechanisms, such as repeat expansion and integration of transposable elements (TEs), can still generate substantial variation^16^. TEs represented a major source of raw genetic material throughout eukaryotic evolution^17^, contributing both to structural and functional innovations^18,19^. These include enhanced DNA processing capabilities such as somatic hypermutation^20^ and V(D)J recombination^21,22^, fusions to transposon-derived domains resulting in alterations of endogenous proteins^23^, and the establishment of regulatory sequences such as centromeres^24^, enhancers^25^, and insulators^26–28^. At the same time, an uncontrolled transposon invasion and proliferation is among the most severe threats to genome integrity^17,29^. Sophisticated defense mechanisms like the PIWI–interacting RNA (piRNA) pathway^30,31^ mediate silencing of TEs through degradation of TE transcripts and preventing their transcription altogether^32^. Genome surveillance is taken a step further in ciliates, such as *Paramecium*^33^, *Tetrahymena*^34^, and *Oxytricha*^35^. Here, TEs and genomic repeats such as minisatellites are permanently excised from the somatic genome as it develops from the germline^36–38^. During post-zygotic development of the progeny’s somatic genome, an intricate mechanism uses small RNAs to compare the contents of the parental somatic and germline genomes^34,38,39^. The selected small RNAs then control transposases^40–43^ for the elimination of any germline-limited sequences that are absent in the parental somatic genome. Consequently, the retention of any sequence in the new somatic genome requires their present in both the maternal germline and somatic genomes^38^. Curiously, although this system enables an efficient and precise elimination of TEs and repeats, it only acts on the developing somatic nucleus and does not eliminate sequences from the germline genome. This is paradoxical, as the persistence of germline limited sequences that must be de-novo eliminated with each generation appears unnecessary, especially since the tools for their removal are available. Considering that the process of programmed DNA elimination in ciliates is evolutionarily ancient^44,45^, we hypothesize that these germline-limited sequences do serve functional roles. These could include a function as a regulatory layer for nuclear development-specific gene expression, or providing feedback checkpoints for the process, or acting as a reservoir for genomic variability that facilitates genomic evolution through emergence or modification of the cellular proteome.

## Results

### Assembling the germline genome of *P.tetraurelia*

Considering the presence of an evolutionary ancient, precise and efficient mechanism of elimination of repetitive and TE-derived sequences from the somatic genome in *P. tetraurelia*, it is puzzling that these sequences persist in the germline. We hypothesized that the germline serves functions beyond simple transgenerational storage of genetic information, and we set out to identify them. An initial obstacle is the absence of a contiguous germline assembly, since the best available draft is fragmented into 26,186 contigs. The complexity of this undertaking arises from the disproportionally higher ploidy of the somatic genome, combined with nearly identical sequence content, except for the eliminated sequences. Thus, to improve the germline assembly, we first sought to develop a method for selective enrichment of germline nuclei.

To achieve this, we developed a modified sucrose cushion gradient fractionation approach, commonly used for density-separation of different nuclear types in related organisms. Previous purification attempts in our laboratory suggested that the somatic and germline nuclei may be physically connected. To achieve their separation, we pre-incubated the cells in phosphate-buffered saline solution (PBS) with 0.05 M EDTA. Following a gentle mechanical disruption with a dounce homogenizer, the resulting partial cell lysates were subjected to sequential centrifugation steps designed to progressively deplete the lysate of somatic nuclei (Figure 1a). Microscopic examination confirmed effective nuclear separation and showed that the final fraction was enriched for round DNA-containing organelles consistent in size and morphology with germline nuclei (Figure 1b).

**Figure 1.**
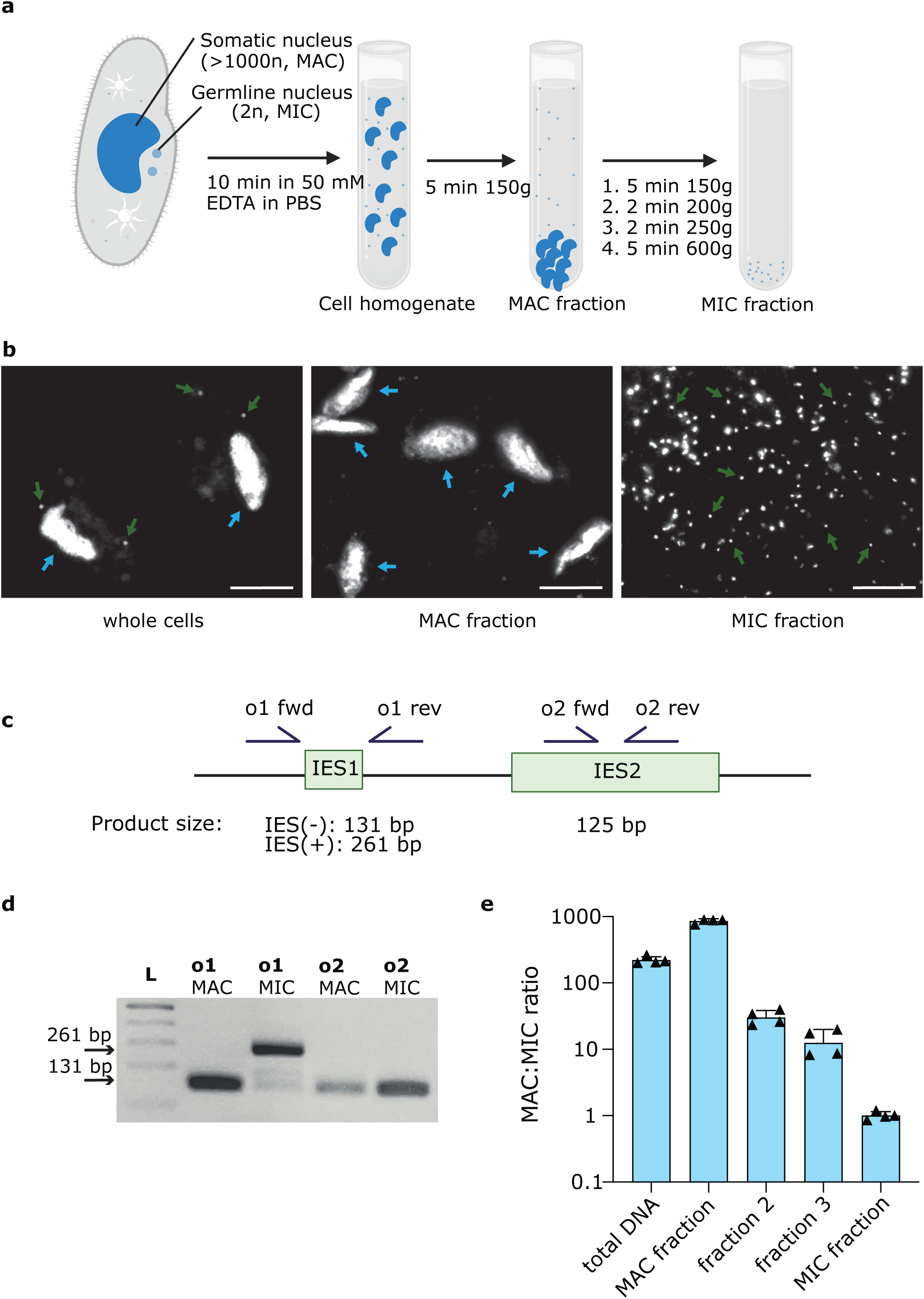
Rapid enrichment of germline nuclei. **a** Schematic of the sucrose-cushion/step-gradient centrifugation used for enrichment of the germline nuclei. **b** DAPI staining of intact cells and of the first and last gradient fractions. Scale bar, 20 μm. Blue arrows indicate somatic nuclei; green arrows indicate germline nuclei. **c** Primer design for enrichment tests: One primer pair flanks a germline-specific sequence (IES1) yielding two product sizes depending on presence/absence of the IES; a second pair amplifies within a germline-limited sequence (IES2). **d** 1% (v/w) agarose gel analysis of PCR products obtained with o1 and o2 primer sets (from panel c) using total DNA (MAC) and germline-enriched DNA (MIC). **e** qPCR analysis of the germline enrichment using the primers shown in panel c, bar plot shows average with error bars indicating SD.

The DNA from the final fraction was tested with two primer sets. One targets somatic genome sequences flanking a short internally eliminated sequence (IES), and the other amplifies an internal region of a germline-limited sequence (Figure 1c). For the IES-flanking primers, PCR yielded a short product in the somatic nucleus fraction, consistent with IES excision, and a larger product in the germline-enriched fraction, consistent with the IES being retained between the primer sites (Figure 1d). Using the same primer sets across the gradient fractions, we observed a progressive increase in the germline-specific products, confirming effective enrichment and validating the efficiency of the separation protocol (Figure 1e). To resolve repetitive and complex genomic regions and exclude potential contaminants (e.g. bacterial DNA), the germline-enriched DNA was subsequently subjected to long-read sequencing.

### The new germline assembly faithfully represents the contents of the germline genome

To remove contaminating reads, we applied an initial filtering step based on a GC-content cutoff of 0.4 (40%). Given that the genome-wide GC content of *Paramecium tetraurelia* is ~27%, this threshold encompassed the entire first peak of read GC distribution (Supplementary Figure 1a). The majority of the filtered reads used for assembly exceeded 3 kb in length (Supplementary Figure 1b), ensuring robust long-read coverage. We employed a haplotype-resolving assembly strategy to retain all eliminated sequences and subsequently selected the longest sequence variants for each genomic region. After additional manual curation and reference-based validation using the previously reported somatic chromosomes, we reduced the number of contigs from 26,186, in the previously published germline genome assembly, to 179 (Table 1). Notably, most contigs were long (N90 = 430,900 bp), and the 132 largest together covered 90% of the assembled genome.

**Table 1.** Summary of the new germline assembly compared with previously published germline and somatic assemblies.

|  | old germline | new germline | somatic DNA | somatic DNA<br>with IESs |
| --- | --- | --- | --- | --- |
| Contig count | 26 186 | 179 | 697 | 697 |
| Largest contig | 286 327 | 1 655 960 | 980 760 | 1 021 975 |
| Total length | 95 031 581 | 101 856 490 | 72 102 941 | 75 658 061 |
| GC (%) | 27.35 | 27.32 | 28.04 | 27.69 |
| N50 | 39 394 | 650 011 | 413 026 | 431 324 |
| N90 | 5 099 | 430 900 | 215 697 | 228 254 |

Next, to assess the completeness of the germline assembly, we used the small RNAs produced from the germline genome during sexual reproduction. During meiosis of the micronuclei, the germline genome is bidirectionally transcribed and gives rise to 25-nt long small, PIWI-bound RNA class, termed scan RNAs (scnRNAs). Upon negative selection of somatic sequences these RNAs guide the programmed DNA elimination of germline sequences in the new somatic nucleus (Supplementary Figure 2a). Therefore, a complete and accurate germline assembly should provide mapping sites for all scnRNAs. When we mapped the 25-nt RNAs obtained from the early developmental stage (20-30% fragmentation), we observed a homogeneous distribution across contigs, with RNA counts scaling proportionally to contig lengths (Supplementary Figure 2b), indicating unbiased genome-wide distribution.

Additionally, a majority of scnRNAs possess a strong 5′ UNG sequence bias (Figure 2a), which can be used to assess the depletion of scnRNAs within the unmapped fraction. Accordingly, the 25-nt RNAs that failed to map to either the previous or the new germline assembly displayed a significantly weaker 5′ consensus than RNAs unmapped to the somatic genome (Figure 2a). Importantly, when the scnRNAs that were unmapped to previous germline assembly were remapped to the new assembly, the characteristic 5′ consensus disappeared entirely (Figure 2a). Consistent with this, while 37% of the 25-nt reads did not map to the somatic genome, fewer than 1% failed to map to either the old or the new germline assembly (Figure 2b). Further investigation revealed that many of the remaining unmapped small RNAs correspond to mitochondrial genome sequences or to an ampicillin-resistance cassette that likely integrated into the genome during earlier laboratory cultivation (Figure 2c).

**Figure 2.**
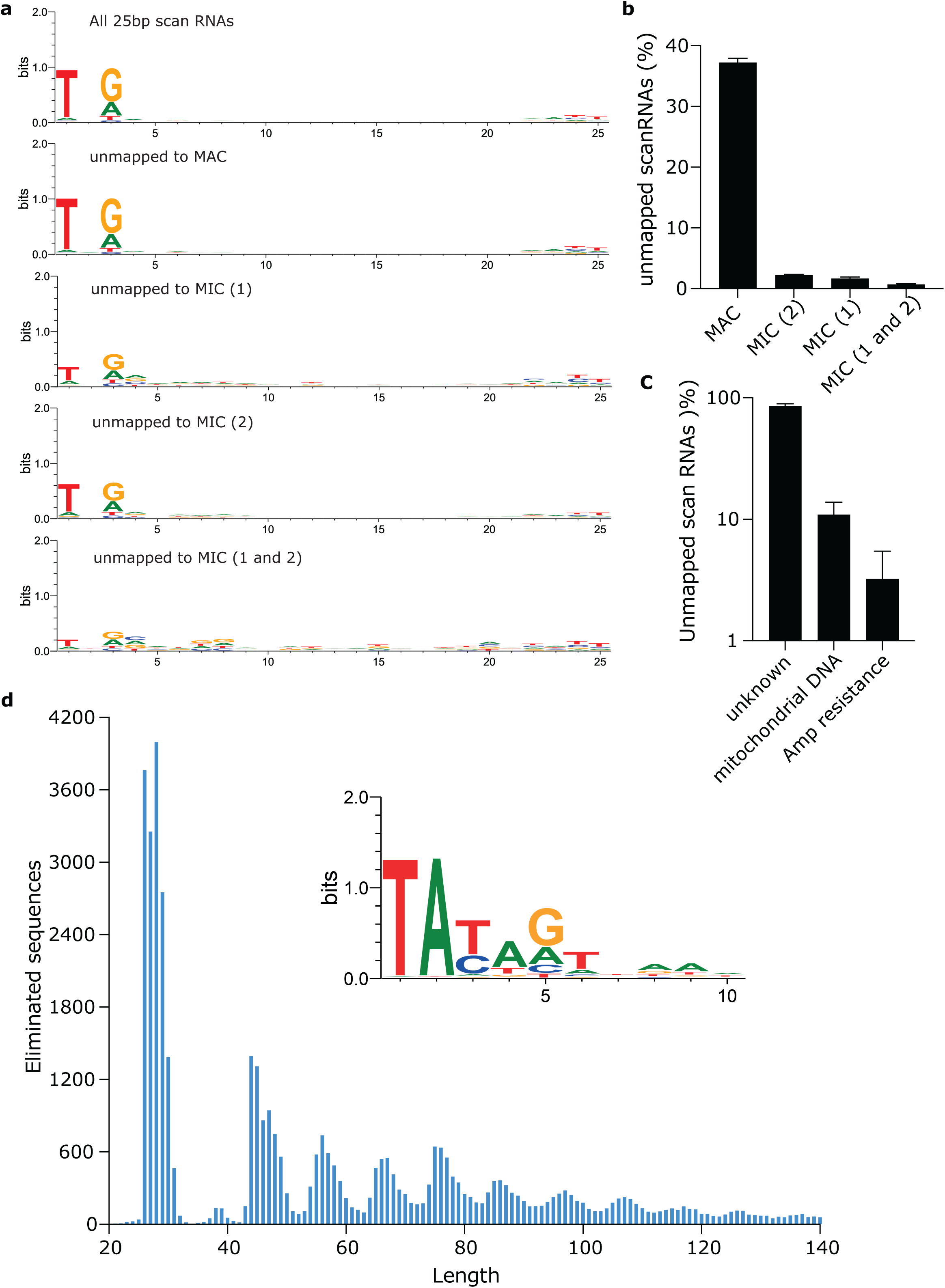
The new germline assembly accurately represents the overall genome content. **a** WebLogo plots showing sequence conservation for all 25-nt scnRNAs, and for the subsets unmapped to the somatic (MAC), previous germline (MIC 1), new germline (MIC 2), and unmapped to both germline assemblies. **b** Bar-plot indicating the percentage of 25-nt scnRNAs from the total 25bp population without full-length matches to the indicated assemblies. Three biologically independent samples were used in small RNA sequencing experiments and mapped to the genome assemblies; error bars indicate SD. **c** Unmapped scnRNAs from panel b analyzed for full length matches to mitochondrial DNA or the ampicillin resistance gene. Bars show three biological replicates and error bars indicate SD. **d** Histogram of IESs smaller than 140 bp identified in the new assembly and WebLogo sequence motif of the 5’ end of these eliminated sequences adjusted for a GC content of 28%.

Another characteristic feature of the germline genome is the presence of tens of thousands of short insertion sequences that are localized within the boundaries of the somatic chromosomes and are therefore named internally eliminated sequences (IESs). To identify insertion/IES segments, we aligned the somatic chromosomes to the new germline assembly using splice-aware settings and then extracted the alignment gaps reported as “introns”. Overall, we identified 46,210 IES sequences, exceeding previous counts, with the largest gains among longer elements (Supplementary Figure 3, Supplementary table 1). Short IESs showed the expected size periodicity consistent with a single helical turn of DNA and lacked any conserved recognition motif beyond the first few nucleotides (Figure 2d). The new germline assembly also allowed us to extend many previously reported IESs that contained secondary internal insertions (Supplementary Figure 4a) as well as those with multiple base-pair substitutions (Supplementary Figure 4b). Together, these results indicate that our assembly provides a nearly complete and accurate representation of the germline genomic content of *Paramecium tetraurelia*.

### Germline-limited sequences frequently produce protein-coding RNAs

Having established that the new assembly accurately represents the germline genomic content, we examined it for the presence of telomeric repeats. Surprisingly, only a few of the contigs carried terminal telomeric repeats. In the somatic nucleus of *Paramecium tetraurelia*, telomeric sequences consist of variable G_(3-4)_C_(2-3)_ hexanucleotide repeats. We therefore considered that this variability might have prevented the incorporation of telomeres by the assembly algorithm. To assess this, we manually screened all reads for the repeat motifs, but out of more than 5 million reads, only 342 contained telomeric repeats (Supplementary data 1). When mapped, the non-telomeric parts of these reads aligned within the interiors of germline contigs, often near eliminated sequences, or to regions near the ends of somatic chromosomes, rather than to terminal positions of germline contigs. The low abundance, combined with their localization, suggest that these reads likely originated from minor contamination with somatic DNA and do not represent genuine germline telomeric sequences.

This unexpected scarcity of telomeric repeats is in strong contrast with the sequence coverage for the remaining parts of the germline genome. Notably, the mean coverage across the assembly was high (216x), and uniform for most contigs (Supplementary Figure 5). Furthermore, the long-read sequencing technology was previously used for detecting telomeric and other complex repeat structures in diverse organisms making a technical fault unlikely ^46,47^. This concludes that either the telomeric sequences were lost during the germline DNA extraction procedure, or that the *Paramecium* germline employs a mechanism that is not based on canonical telomerase. Such system is known in *Drosophila*, where the chromosomal termini are extended through replication of non-LTR retrotransposons. We therefor examined the terminal 10kb regions of the germline contigs.

Remarkably, we detected a pair of inverted repeats in both ends of contig 2 (Figure 3a). To test whether these terminal sequences are transcriptionally active, we performed full-length poly(A)+ RNA sequencing and detected transcript isoforms that mapped precisely to these inverted repeat regions (Figure 3a). Close inspection revealed an uninterrupted open reading frame spanning nearly the entire transcript. The four repeat-encoded proteins on contig 2 show extremely high sequence homology (Figure 3b) and WD40 repeats, consistent with a conserved origin and potentially even functional roles, likely in protecting chromosomal ends or scaffolding of protein complexes at the telomeric regions, as observed in other organisms^48,49^. Overall, we discovered that this conserved sequence is present at both ends of 63 contigs and at least one end of 137 out of the 179 contigs (Figure 3c). Further work is needed to determine whether these germline-limited DNA repeats, their transcripts, or the encoded proteins participate in a non-telomeric mechanism for chromosomal end maintenance, or if they have other roles for the chromosomal organization. Notably, other regulatory loci such as the centromeric sequences are also eliminated from the germline during the development of the somatic nucleus.

**Figure 3.**
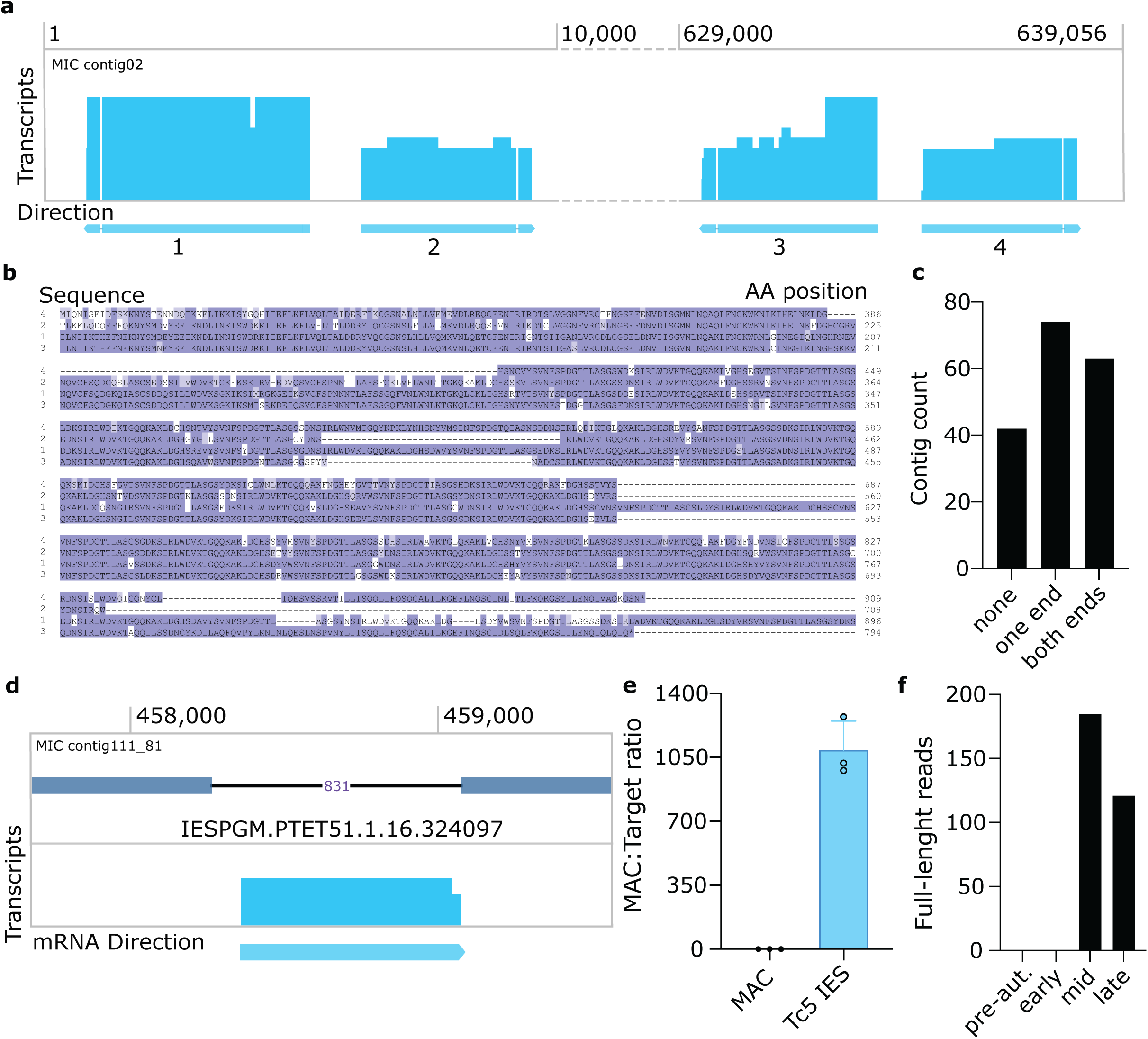
Germline-limited sequences at contig ends are transcribed and encode uninterrupted homologous open reading frames. **a** Schematic diagram of 10 000 bp at the 5’ and 3’ ends of contig_02 showing highly homologous, transcribed, and spliced inverted repeats. **b** The four terminal transcripts from panel a encode uninterrupted homologous open reading frames containing WD40 domains. **c** Histogram of contig counts with at least one or both termini (within 10 kb) containing WD40-encoding repeats. **d** Schematic showing the germline-limited internal eliminated sequence IESPGM.PTET51.1.16.324097 from which polyadenylated transcripts are produced. **e** qPCR validation of the relative abundance of the Tc5-derived sequence (shown in panel d) compared with the somatic genome. Error bars indicate SD. **f** Counts of full-length transcript reads matching the Tc5 germline-limited transposon across different developmental stages.

Importantly, these results demonstrate that at least some germline sequences are actively transcribed into protein-coding mRNAs. This challenges the prevailing view of the germline as a completely inert repository of genetic information. As a follow-up, we asked whether other germline-limited regions are also transcribed, at least during the sexual cycle. Full-length RNA sequencing revealed thousands of transcript isoforms originating from germline-specific loci (Supplementary Figure 6). Notably, these transcripts were detected shortly before and, to a larger extent, during sexual reproduction. The detected transcripts arose from two major sources: TE sequences and repeats flanking the ends of somatic chromosomes, as well as from longer IESs embedded within somatic chromosomes. We manually verified over 100 long eliminated sequences that contribute to transcripts during the formation of the new somatic nucleus (Supplementary data 2). Among these, one eliminated sequence (PDB, IESPGM.PTET51.1.16.324097; 831 bp) caught our attention due to high levels of transcription (Figure 3d). It encodes a putative DNA-binding domain homologous to that of Tc5 transposons, supporting a transposon-derived origin for this IES. Furthermore, the gene adjacent to this germline-limited sequence shows much weaker expression than the transposon itself. We further confirmed via qPCR that the DNA itself is eliminated from the somatic genome (Figure 3e). The expression of the transcript is stage-dependent (Figure 3f), and the encoded protein shows high amino-acid similarity to several Tc5 transposon-derived proteins that are retained and expressed from the somatic genome (Supplementary Figure 7). This observation combined with the high expression level suggests that a co-option process for these Tc5 transposon-derived DNA binding domains is currently underway. While most of the homologous sequences have already been domesticated and are retained in the somatic genome, the germline Tc5 element can only be transiently expressed during the development of the somatic nucleus.

### Short transposon remnants contribute to transcriptome complexity

Encouraged by the observed transcriptional activity of long germline-limited sequences, we next asked whether shorter transposon remnants also contribute to the cells’ transcriptome. Comparison of the eliminated-sequence set with all detected transcript isoforms revealed thousands of isoforms containing short insertions derived from eliminated regions. We then focused on the shortest, highly abundant class of eliminated sequences (26–31 bp) (Figure 4a), whose length is similar to *Paramecium* introns. For each length, only a subset of eliminated sequences appeared in the transcriptome, but the proportion incorporated into transcripts varied with sequence length (Figure 4a). Notably, the most likely insertion sizes were 27 bp and 30 bp, both multiples of three, implying a translational constraint favoring in-frame insertions (Figure 4b). Previous results demonstrated that the nonsense-mediated decay (NMD) pathway is strongly upregulated in *P. tetraurelia*, suggesting that transcripts containing premature stop codons are rapidly degraded^50^. To test whether the observed length bias reflects selective pressure against frame-disrupting insertions, we examined the resulting proteome for proteins that were altered due to the insertion of an IES in their transcripts (Figure 4c, Supplementary Table 2). Indeed, most proteins with longer than wild-type sequences were derived from transcripts containing 27bp or 30bp insertions (Figure 4c). In contrast, insertions of other lengths commonly generated frameshifted, truncated proteins (Figure 4c, Supplementary Figure 8a). Still, the presence of only one termination codon allowed many such frameshifts to produce proteins of similar or even greater length than wild type (Figure 4c). We also found that in-frame stop codons are slightly underrepresented for the insertions of 27bp and 30bp lengths (Supplementary Figure 8b). Consequently, in most cases, these 27bp and 30bp insertions introduced short peptide extensions within existing domains, while leaving the rest of the protein sequence unchanged as demonstrated for an autogamy-specific DNA-ligase (PBD, PTET.51.1.P0010027) (Figure 4d). The expression levels of modified and unmodified transcripts showed stage dependency but were overall comparable for this gene (Figure 4e). This suggests a potential functional role, such as linking the functionality of proteins with the progression of DNA elimination from the genome. These inserted sequences could therefor represent an additional layer of gene regulation, consistent with our previous report that IES insertions in the promoter regions regulate gene expression^51^.

**Figure 4.**
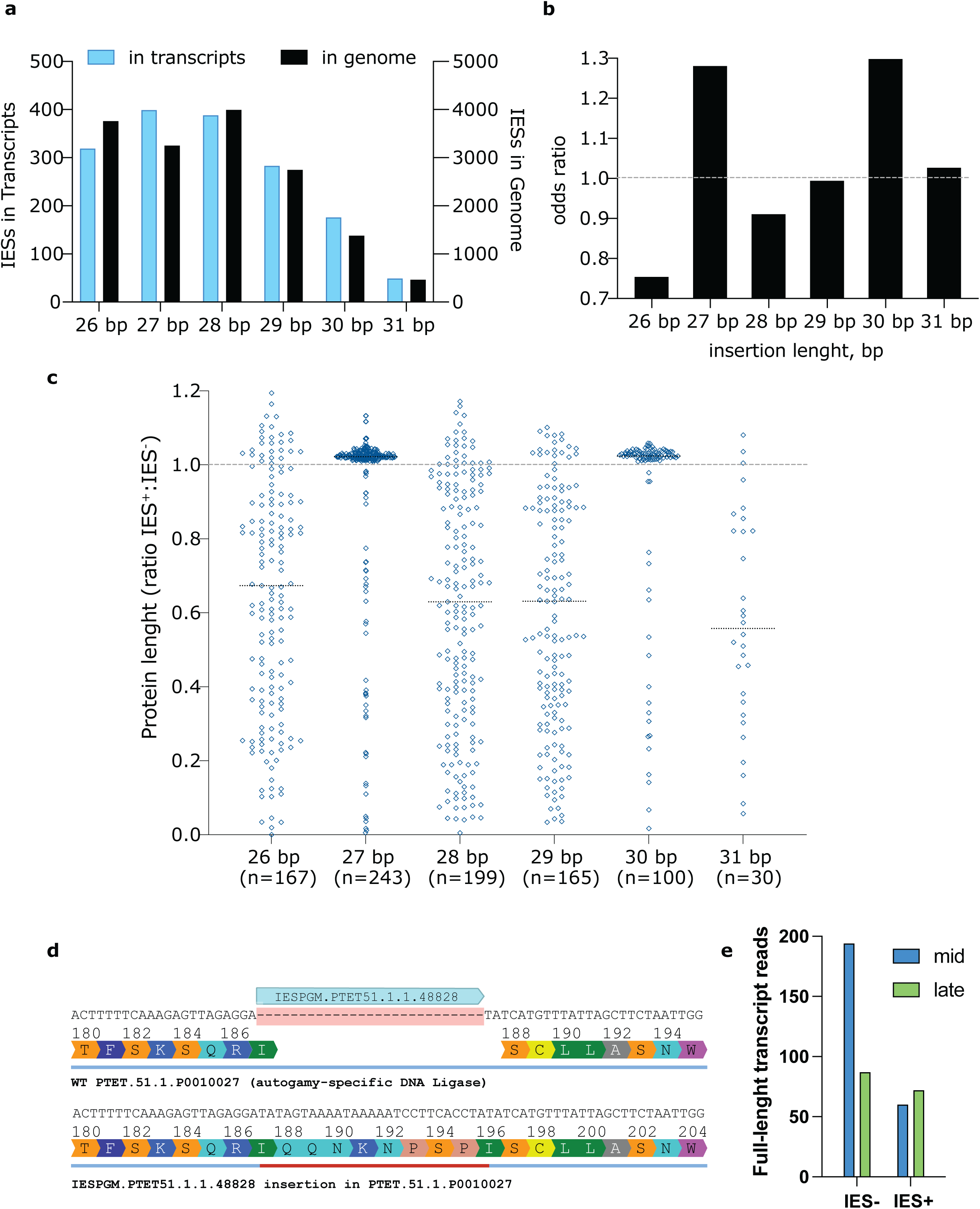
Short transposon remnants generate transcriptome diversity. **a** Comparison of 26 to 31 bp transposon remnants (IESs) present in genomic DNA versus those found in polyadenylated transcripts. **b** Odds ratio for each transposon remnant size in the range 26 to 31bp, calculated as the fraction in the genome divided by the fraction observed in the transcripts. **c** Effects on open reading frames caused by insertion sequences. Each point is the ratio of the modified to unmodified ORF for a single transcript, median value per group is shown as a dotted line. **d** Transcripts are generated from DNA in which the transposon remnant IESPGM.PTET51.1.1.48828 is excised, as well as from DNA in which it is retained. The retention results in a 9 amino acid extension of the protein without other alterations to the amino acid sequence. **e** Counts of full-length transcript reads with or without the insertion sequence IESPGM.PTET51.1.1.48828 from mid (50%) and late (100%) developmental stages.

Collectively, our results show that short transposon remnants contribute to subtle, yet widespread diversification of the transcriptome and, by extension, the proteome. In the highly polyploid somatic nucleus, rare IES retention events can co-exist with unmodified alleles. Upon selective pressure from changing environments or fitness improvements the populations enriched in modified variants could outcompete others. Because DNA elimination is in large proportion guided by the scnRNA pathway (Supplementary Figure 2a), sequences retained in one generation are more likely to be retained in the next, creating a positive feedback loop (Supplementary Figure 9). The co-existence of a somatic and germline nuclei with different genomic contents could therefor provide a potential mechanism for driving incremental protein modifications and thereby promote genome evolution.

## Discussion

In this study, we established the first nearly complete germline genome assembly of *Paramecium tetraurelia*, reducing the number of contigs from 26,186 to only 179. This represents a substantial improvement in continuity and completeness. Our selective enrichment protocol for isolating germline nuclei is simple and reproducible, requiring neither fluorescence-activated cell sorting nor transgenic labeling, unlike previous approaches^52^. The final contig count corresponds well to the estimated count of germline chromosomes^53,54^, providing further confidence in the assembly.

Additionally, because scnRNAs are generated from the entire germline genome^38^, the mapping of nearly all scnRNA to our assembly strongly supports its completeness. Interestingly, we also detected scnRNAs mapping to specific regions of the mitochondrial genome, which to our knowledge has not been reported before. These scnRNAs are not distributed across the whole mitochondrial sequence but cluster within defined regions, suggesting that mitochondrial genome fragments may have overlapping sequences with the germline genome, or could participate in a crosstalk between nuclear and mitochondrial compartments.

The unexpected scarcity of telomeric reads in the germline assembly contrasts with expectations, as the same sequencing technology reliably retrieves telomeric repeats in other organisms. While it impossible to exclude that the DNA enrichment procedure selectively discarded telomeric sequences, an alternative explanation could be a non-canonical mode of chromosome-end replication. The consistent presence of highly homologous transcribed repeat sequences near the ends of most contigs supports this notion. In Drosophila, the telomeres are maintained through transposon-based mechanisms rather than canonical repeats^55,56^, and furthermore, WD40-domain scaffolding proteins as those encoded by the end repeats, are known components of telomere-processing complexes in other eukaryotes^48,49,57^. Additional studies are needed to uncover if the WD40 end repeats that we identified are involved in any form of chromosomal end replication, including subtelomeric or telomere-replication pathway.

Our results also provide direct evidence that long germline-limited sequences, including those internal to somatic chromosomes, are actively transcribed. This finding challenges the long-standing assumption that the germline nucleus serves merely as a transcriptionally inert genetic archive. The detection of germline transcripts across multiple developmental stages indicates that some are expressed early, whereas others accumulate later during autogamy, when the new somatic nucleus forms. These patterns suggest that transposon-derived elements may differ in their timing and mechanism of expression, potentially reflecting distinct stages of domestication or regulatory specialization^17^.

At a conceptual level, our findings also expand the functional interpretation of the nonsense-mediated decay (NMD) pathway. Rather than acting solely as a quality-control mechanism, NMD may facilitate the positive selection of modified transcripts by preferentially stabilizing variants with preserved reading-frame, thereby fostering evolution. This is particularly relevant in *Paramecium*, where the natural base-substitution rate is extremely low^12^, implying a reduced role for point-mutation–driven evolution. Additionally, the genetic code of ciliates is non-standard, often having only one stop codon^58^, or even no dedicated stop codon at all^59^. The highly specific translation machinery ensures the correct expression, even of frameshifted sequences^58^, which can further increase the tolerability for discreate modifications of protein domains.

Protein fusions^23^ and truncations^60^ are longstanding sources of genetic innovation and are also routinely exploited in the laboratory to engineer proteins with desired functions^61^. Such engineered proteins hold the potential for tremendous improvements of biological pathways^62^ and are also randomly occurring at a natural setting^63^. The main disadvantage of this process comes with the fundamental fitness cost of change to the proteome of an organism, as most modifications are more likely to be deleterious^5^. In this context, the combination of nuclear dimorphism and programmed DNA rearrangements may therefore represent an elaborate system for generating controlled genomic variability through insertion and excision events (Supplementary Figure 8). In this framework, the germline serves not only as a barrier protecting against transposable elements but also as source for evolutionary innovation^16,38^, allowing transient probing of new sequence combinations without compromising somatic genome integrity at large. The presence of short insertion sequences such as transposon remnants^42^ within coding and regulatory regions supports this model. Some insertions subtly alter protein termini or modulate promoter activity, while excision restores canonical forms. Such reversible genomic plasticity could provide an additional regulatory layer, coupling genome rearrangement to stage-specific gene expression. Although the changes described here occur in the somatic genome, its phenotypes are among the primary drivers of germline genome evolution, because selection acts on somatic genome traits. These will, in turn, affect the somatic genome traits of the offspring via an RNA-mediated nuclear crosstalk^38^. Therefore, germline alleles that lead to improved somatic fitness will leave more descendants, which can shift the germline allele frequencies across generations. Together, our results suggest that *P. tetraurelia* has likely evolved a dual-purpose nuclear dimorphism: on one side as a safeguard for genomic stability, while additionally fostering a controlled diversification of the transcriptome and proteome.

## Methods

### Cultivation

Monoclonal wild type *Paramecium tetraurelia* strain 51 cell lines were cultivated as described previously ^64^. In brief, a single cell was transferred to a depression slide well containing wheat grass powder (Pines International) light medium (0.2x) supplemented with 0.8 μg/mL β-sitosterol (Sigma-Aldrich, cat. no. 567152) and bacterized with *Klebsiella pneumoniae*. The cell density was maintained below 500 cells/mL, to prevent premature initiation of the meiotic reproduction cycle. When 50 vegetative cell divisions were reached, the cells were transferred to bacterized wheat grass powder rich medium (1x) supplemented with 0.8 μg/mL β-sitosterol. The cell density was maintained below 1000 cells/mL with addition of new medium, until a total volume of the culture of 1L was reached. The culture was then further incubated at 27 °C until all bacteria were consumed, and the starvation induced initiation of the sexual reproduction cycle. For extraction of the germline nuclei, 200 mL of the postautogamous culture were added to 1.8L of bacterized wheat grass powder rich medium (1x) supplemented with 0.8 μg/mL β-sitosterol and incubated for two days at 27 °C before harvesting the cells. This ensures that the cells are less than 10 divisions old and cannot initiate the meiotic cycle, thereby preventing the formation of somatic nuclear fragments that could contaminate the germline fraction.

### Germline DNA extraction

To obtain the germline nuclei, the cultures were first filtered through a tissue cloth, and the cells were pelleted for 2 min at 280 g using an oil-measurement centrifuge fitted with pear shaped flasks. The cell pellet was washed with phosphor buffered saline (PBS) pH7.4, and then resuspended in ten volumes of PBS supplemented with 0.05 M EDTA (Fischer Scientific cat. no. AAA151610B) and then incubated for 5-10 minutes. The pellet was washed twice with PBS and then resuspended in three volumes nuclear extraction buffer (250 mM Sucrose, 10 mM MgCl_2_, 0.5% (v/v) Triton X-100, 0.5% (w/v) NP-40). The cell membrane was disrupted with 5-10 strokes in a tight dounce homogenizer, and the progress of the lysis was regularly controlled for fraction of released nuclei using DAPI staining. The lysate was then filled up to 10 mL in a falcon tube, centrifuged 5min at 150g, and the supernatant was transferred to a new tube. The step was repeated using 5 min at 150g, then 2 min at 200g, then 2 min 250g, and after the final transfer of supernatant to a new tube, the germline nuclei were pelleted for 5 min at 600g. The pellet was resuspended in 1 mL nuclear wash buffer (250 mM Sucrose, 10 mM MgCl_2_) then transferred to 1.5 mL Eppendorf tube, and centrifuged for 2 min at 3000g, and after an additional wash step, the pellet was shock-frozen in liquid nitrogen. The DNA was extracted using the NucleoSpin Tissue genomic DNA extraction kit (Macherey-Nagel, cat. no. 740952.50) according to the manufacturers protocol for processing of tissue samples. In short, the pelleted nuclei were incubated with buffer T1 and Proteinase K for 1h at 56 °C, upon addition of buffer B3 the samples were incubated at 70 °C for 10 min, mixed with EtOH and loaded on the silica membrane. After washes with buffers BW and B5, and drying of the membrane, the DNA was eluted in 50 μL of 70 °C preheated elution buffer (5 mM Tris/HCl, pH 8.5).

### Total RNA extraction

800 mL of Paramecium culture in wheat grass powder rich medium supplemented with 0.8 μg/mL β-sitosterol were used for extracting total RNA at each timepoint,. The cultures were filtered through a tissue cloth, and the cells were pelleted for 2 min at 280 g using an oil-measurement centrifuge fitted with pear shaped flasks. The pellets were transferred to 15 mL falcon tubes and washed three times with PBS before shock-freezing in liquid nitrogen. Total RNA was extracted using TRI reagent (Thermo Fischer, cat. no. AM9738) according to the manufacturers protocol. In brief, the pellets were resuspended in three volumes TRI reagent, vortexed vigorously and incubated for 5 min at r.t., then 0.2 mL of chloroform was added for each mL of TRI reagent. Upon vigorous agitation, and incubation for additional 5 min, the samples were centrifugated for 10 min at 21 000 g and 4 degrees °C, and the aqueous phase was collected. Next, the nucleic acids were precipitated by addition of one volume of ice-cold isopropanol, followed by pulse-vortexing and incubation for 20 min at r.t., before centrifugation for 15 min at 21 000 g and 4 degrees °C. The RNA pellets were washed with 75% EtOH, centrifuged for 5 min at 21 000 g and 4 degrees °C, dried and dissolved in ultra-pure H_2_O.

### Quantitative PCR

The ratio of germline DNA to somatic DNA in a sample was quantified using parallel qPCR reactions conducted using KAPA SYBR FAST qPCR 2x Master Mix (Roche, cat. no. KK4601) according to the manufacturers protocol. For each reaction, 1 ng of DNA was used in a total volume of 10 μL with 0.5 μM forward and reverse oligonucleotides. The reaction was conducted on a Bio-Rad CFX384 using skirted 384 well plates and fluorescence measured for the SYBR preset. The cycling program used 95 °C for 20s initial denaturation, followed by 40 cycles of two-step amplification using 4s at 95 °C denaturation and 20s at 60 °C for annealing and elongation.

### PacBio Iong-read sequencing

Long read PacBio sequencing was conducted by the Next Generation Sequencing Platform of the University of Bern according to their standard operational procedures. In brief, for full length RNA sequencing the Kinnex RNA library workflow was followed. First strand synthesis was done using 300 ng of total RNA with RIN 10 using oligo-dT primers and Iso-Seq template switch oligonucleotides. The cDNAs were cleaned up using SMRTbell beads and amplified and barcoded using IsoSeq v2 barcoded cDNA primer. 55 ng of the barcoded cDNAs were used in the Kinex PCR reaction and upon cleanup were ligated to a complete Kinnex array, with SMRTbell adapters attached to the ends. The arrays were sequenced on a PacBio Revio instrument (Software version 13.0.0.212033, Chemistry version 13.0.0.205983) and the data was processed using SMRT Link version 13.1.0.221970. The HiFi reads containing concatenated transcripts were segmented to S-reads using the read segmentation data utility and then processed using the PacBio Iso-Seq workflow to cluster reads to individual full-length transcript isoforms. For sequencing of the genomic DNA, the purified germline DNA was size selected to 5-10 kb post extraction and 20ng of the cleaned-up DNA was used in the PacBio AmpliFi workflow. In brief, the DNA was repaired and A-tailed, then ligated to the Twist universal adapters and PCR amplified using the Twist UID primers by KOD Xtreme Hot Start DNA polymerase. The amplified DNA was repaired and A-tailed and ligated to SMRTbell adapters before sequencing on a PacBio Revio instrument (ICS version 13.3.0.253824).

### Bioinformatic analysis

The segmented PacBio HiFi sequencing reads from the germline DNA were filtered for a GC content lower than 40% and the remaining 5,007,860 reads were used in Hifiasm v0.25.0 for obtaining 357 contigs in the primary pseudohaplotype assembly, assembly quality was then verified with Bandage 2022.09. Additional contigs from the alternate assembly were included based on exact matches to contig ends or germline limited sequences. The contigs were additionally checked for overlaps using minimap2 using the (self-homology) present, and also the present somatic genome assembly (PDB^65^) using the (map-hifi) preset, and then manually curated based on the overlaps so that every somatic chromosome is localized in only one germline contig, thereby reducing the final contig count to 179. The properties of the final assembly were analyzed with Quast v5.3.0 and compared to the previously published genome assemblies. For assessing the genome content, we used previously published small RNA sequencing results (SRR17246034, ERS1658939, ERS1658942) and mapped the 25bp long early stage scnRNAs using bowtie2 v2.5.3 with the (--very sensitive end to end) preset. Sequence consensus for the unmapped scanRNAs were generated using WebLogo 3.7.9 with an adjustment for a 28% GC content. Internal eliminated sequences (IES) in the genome were defined as introns using minimap2 v2.28 with the --splice-flank preset to map the somatic assembly to the new germline assembly. The intron ranges were extracted using bedtools v2.31.1 getfasta. In cases of ambiguity due to start and end symmetry of the sequence, a TA dinucleotide was selected as the starting point. For assessing the consequences of ultrashort TE remnant insertions into transcripts, we used blastx to compare the transcripts against the known Paramecium tetraurelia 51 proteome (PDB) to determine the translation start site. Next, the 5’ UTRs were trimmed to the matching start site located in the first 150 bp of each transcript both for the transcript with the insertion, and its variant without the insertion, translations were generated using getorf for the first reading frame and the lengths of the resulting translation products were compared. The count of in frame stop codons was determined by filtering the translation products with a length corresponding to the location of the ultrashort TE remnant in the transcript. Most calculations were done using resources provided by the European Galaxy server 35446428. Additionally, data was pre- and post-processed in Microsoft excel, data visualizations were generated in Prism GraphPad. Clustal omega was used for multiple sequence alignments and determining phylogenetic relationships.

## Acknowledgements

The authors are thankful to Robin Hogg for helpful discussions and Nasikhat Stahlberger for technical support. The authors are furthermore thankful to the Next Generation Sequencing Platform of the University of Bern and Pamela Nicholson for experimental assistance and advice. The Galaxy server used for some calculations is partly funded by the German Federal Ministry of Education and Research BMBF grant 031 A538A de.NBI-RBC and the Ministry of Science, Research and the Arts Baden-Württemberg (MWK) within the framework of LIBIS/de.NBI Freiburg. A University of Bern Biorender license was used for generating some of the illustrations. This project was supported financially by a Swiss National Science Foundation grant nr. 214853 awarded to M.N and Swiss National Science Foundation grant nr. 229074 and an Initiator grant from the University of Bern awarded to B.-A.S.

## Author contributions

The research was conceptualized by B.-A.S. and M.N., the experiments were conducted and analyzed by B.-A.S. The manuscript was written by B.-A.S. and reviewed by M.N. M.N. and B.-A.S. acquired the funding.

## Data availability

All data is freely available upon request. The full-length RNA sequencing reads, the germline DNA sequencing reads, and the assembled germline genome have been deposited in the European nucleotide archive (ENA) under study accession number PRJEB101472.

---

**Supplementary Figure 1.**
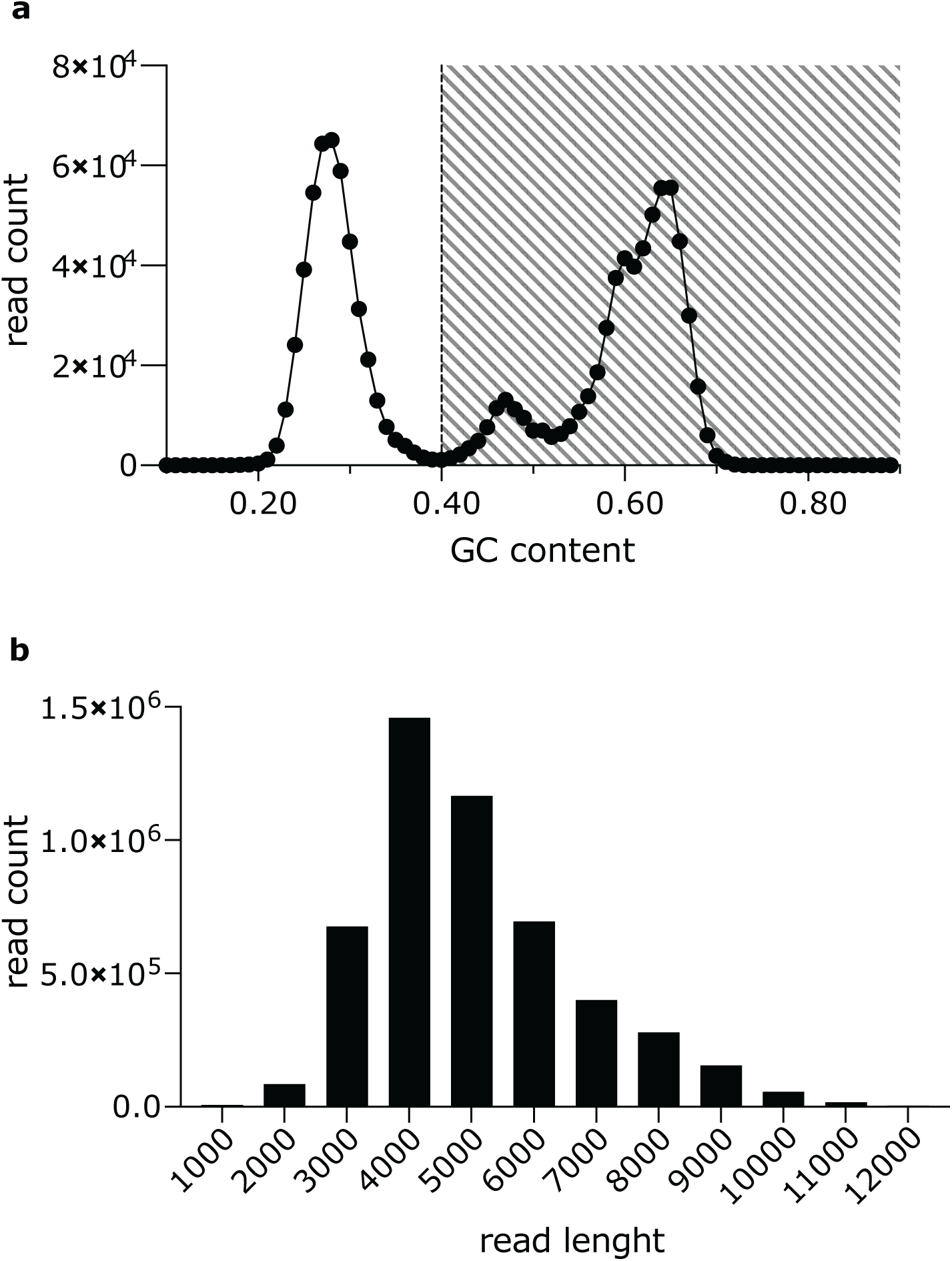
Filtering of long reads for the germline genome assembly. **a** Distribution of per-read GC content for the raw dataset; the dashed area shows the fraction of reads that was removed by GC-content filtering. **b** Read-length distribution of the filtered dataset used for the final germline genome assembly.

**Supplementary Figure 2.**
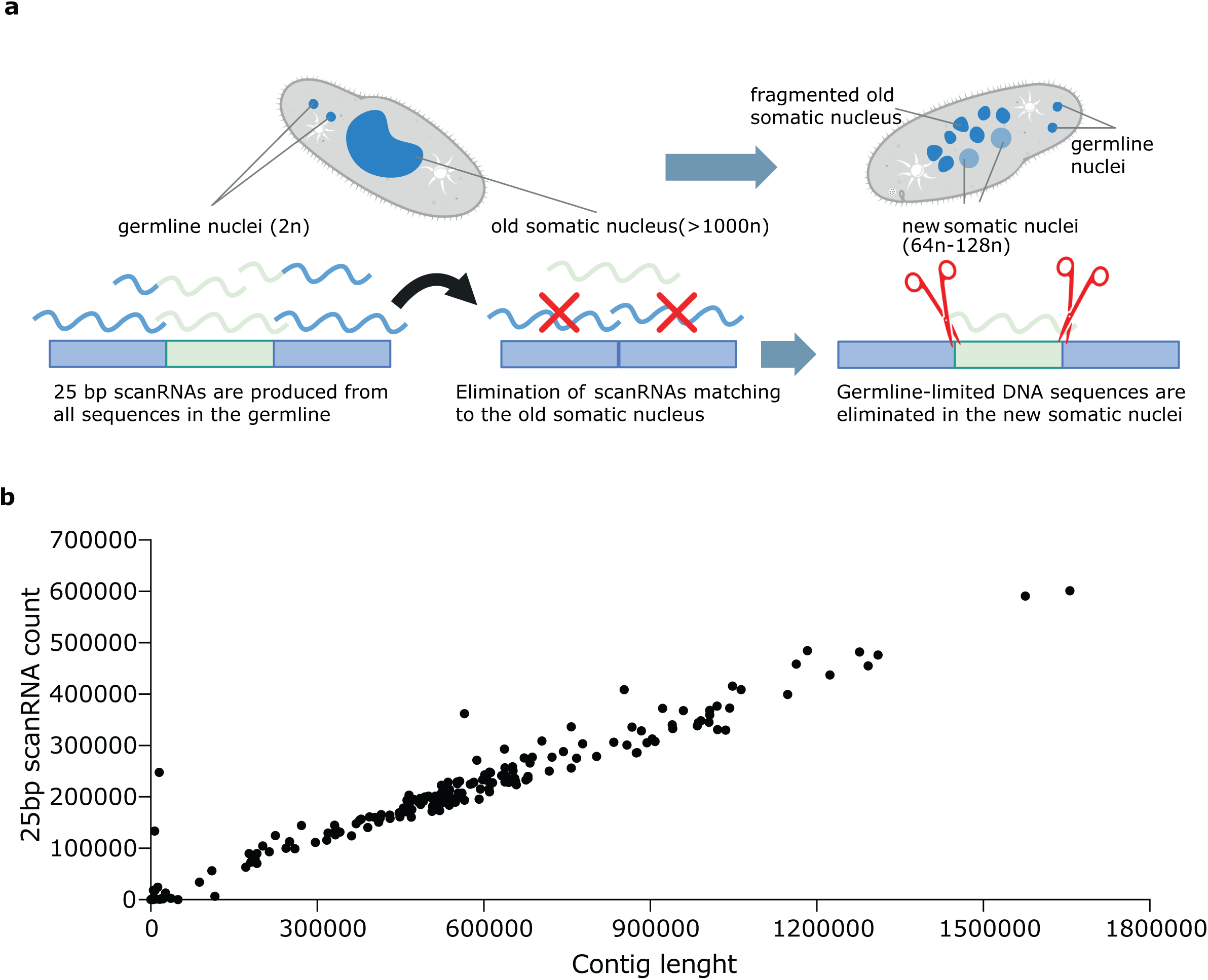
Early stage 25bp scanRNAs map homogenously to the germline assembly. **a** Schematic representation of the scnRNA model in *P.tetraurelia*. 25-nt small RNAs are generated from the entire germline genome. **b** Counts of 25-nt scnRNAs with a full-length matches per contig of the germline assembly.

**Supplementary Figure 3.**
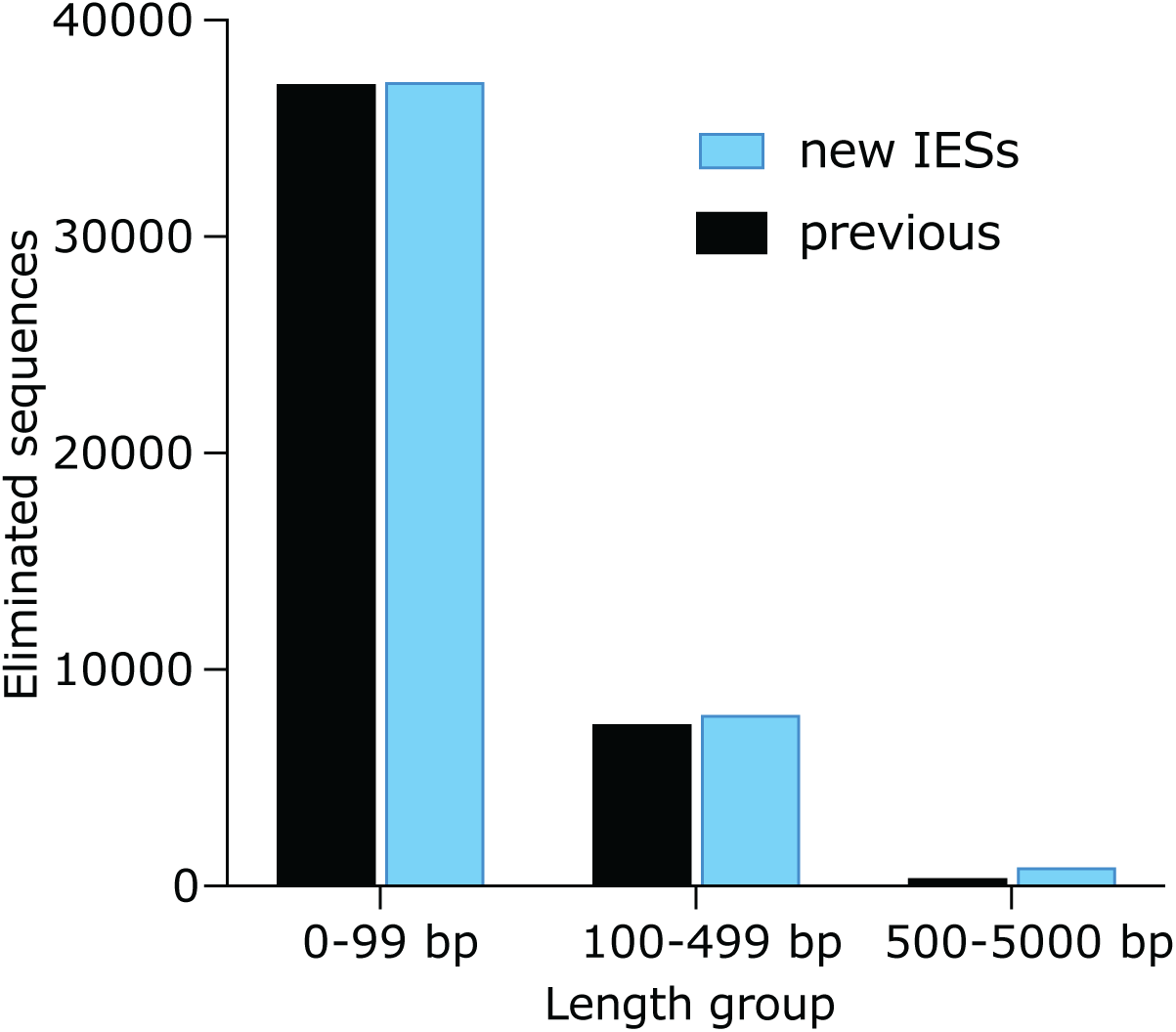
The new germline assembly reveals additional IESs. Histogram plot showing the counts of internal eliminated sequences (IESs) disrupting somatic chromosomes, comparing those previously reported with the newly identified IESs in the present assembly.

**Supplementary Figure 4.**
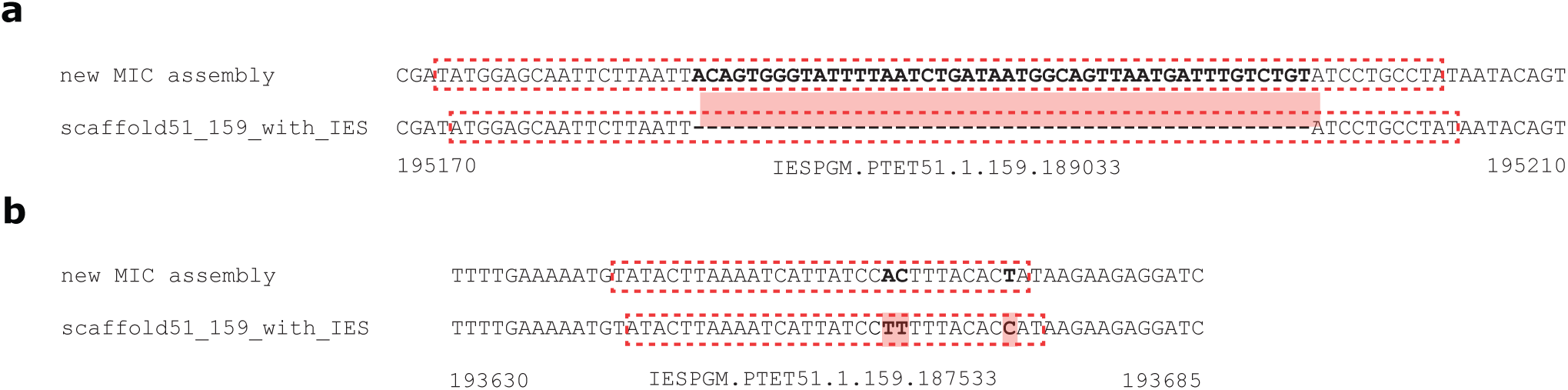
The new germline assembly corrects IES annotations. **a** Example of a previously incomplete IESs, IESPGM.PTET51.1.159.189033, which in the new assembly contains an additional, previously unreported embedded IES sequence. **b** Example of a corrected single-nucleotide error in IESPGM.PTET51.1.159.187533 revealed by the new assembly.

**Supplementary Figure 5.**
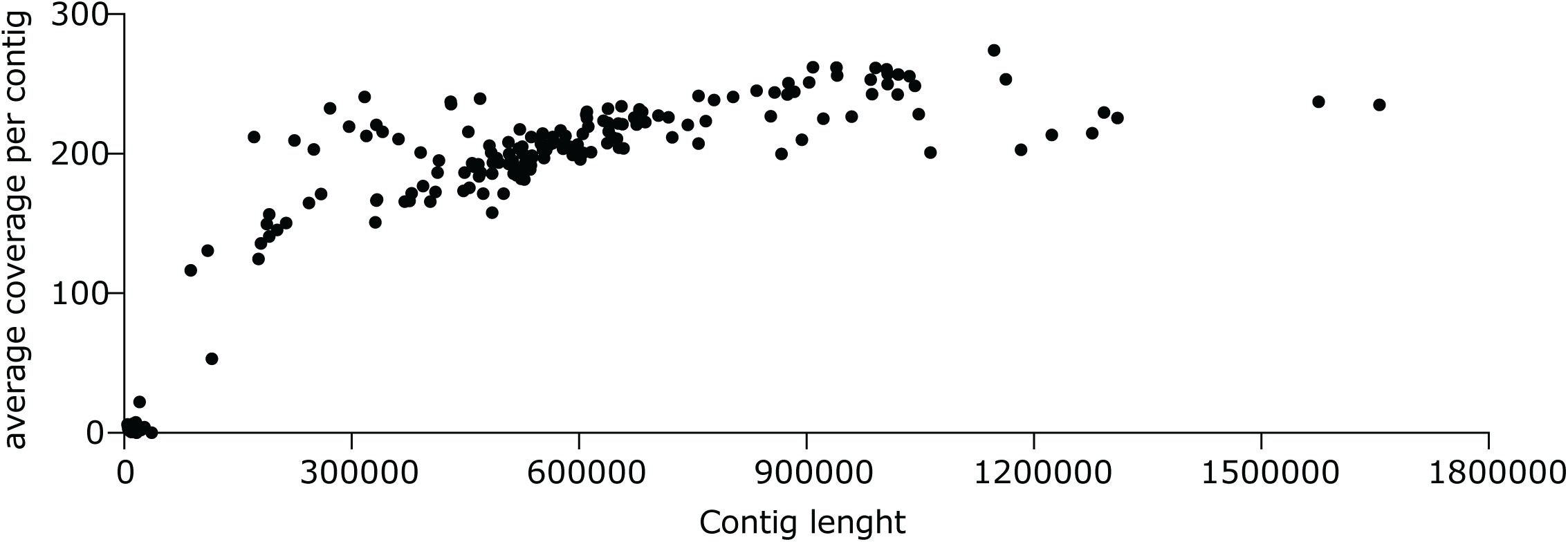
High average read coverage in the majority of contigs. Individual datapoints show the average coverage at each base in the contigs in relation to the length of each contig.

**Supplementary figure 6.**
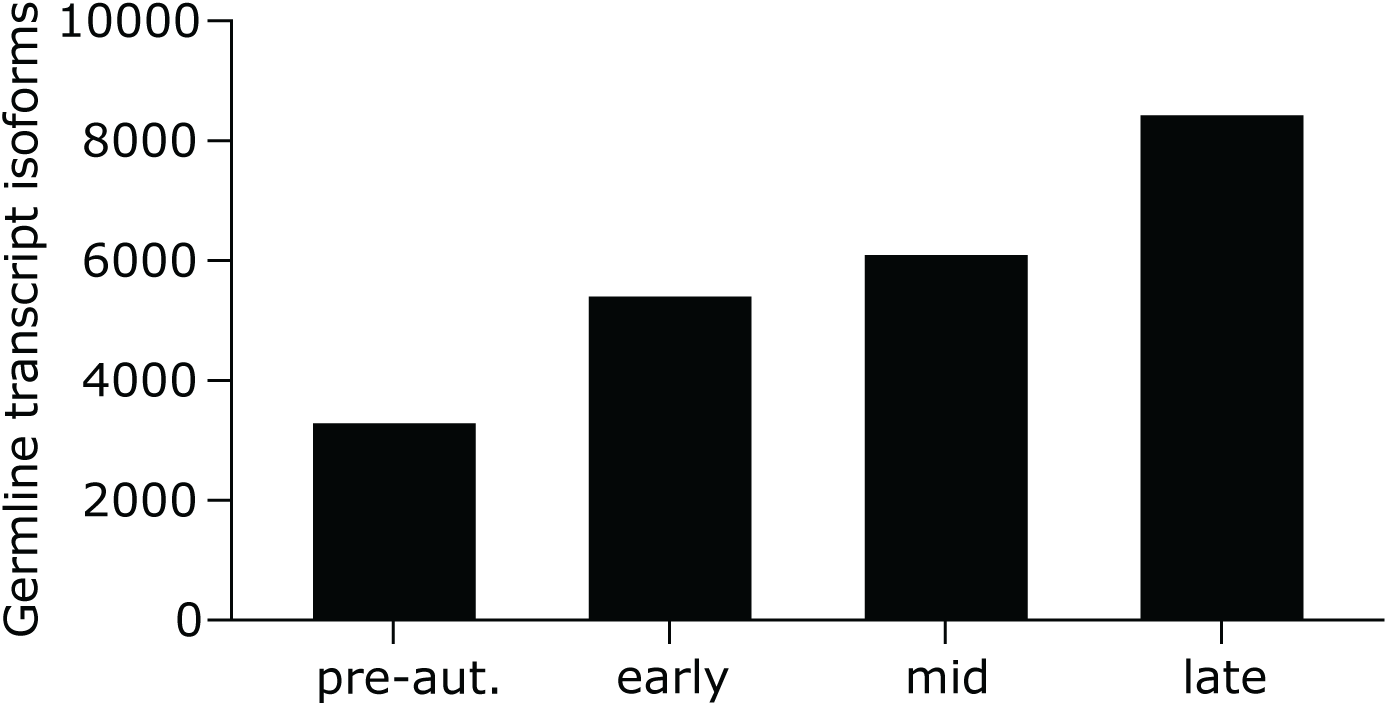
Many transcript isoforms originate from germline limited sequences. Histogram plot of the counts of transcript isoforms from different developmental stages (pre-aut.: vegetative growth just prior to autogamy, early: 10% fragmentation, mid: 50% fragmentation, late: 100% fragmentation) with a match to a germline limited sequence.

**Supplementary figure 7.**
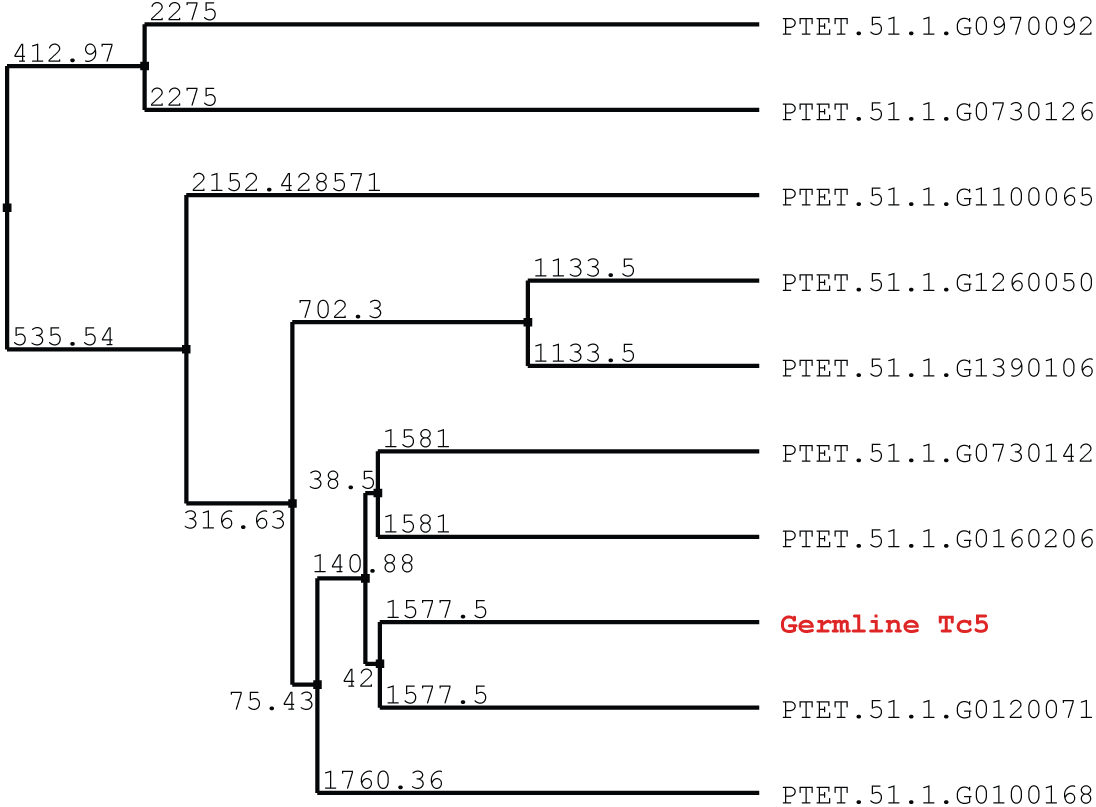
Many transcript isoforms originate from germline limited sequences. Phylogenetic tree of the germline limited Tc5 element encoded by IESPGM.PTET51.1.16.324097 showing its homology to multiple protein-coding sequences retained in the somatic genome.

**Supplementary Figure 8.**
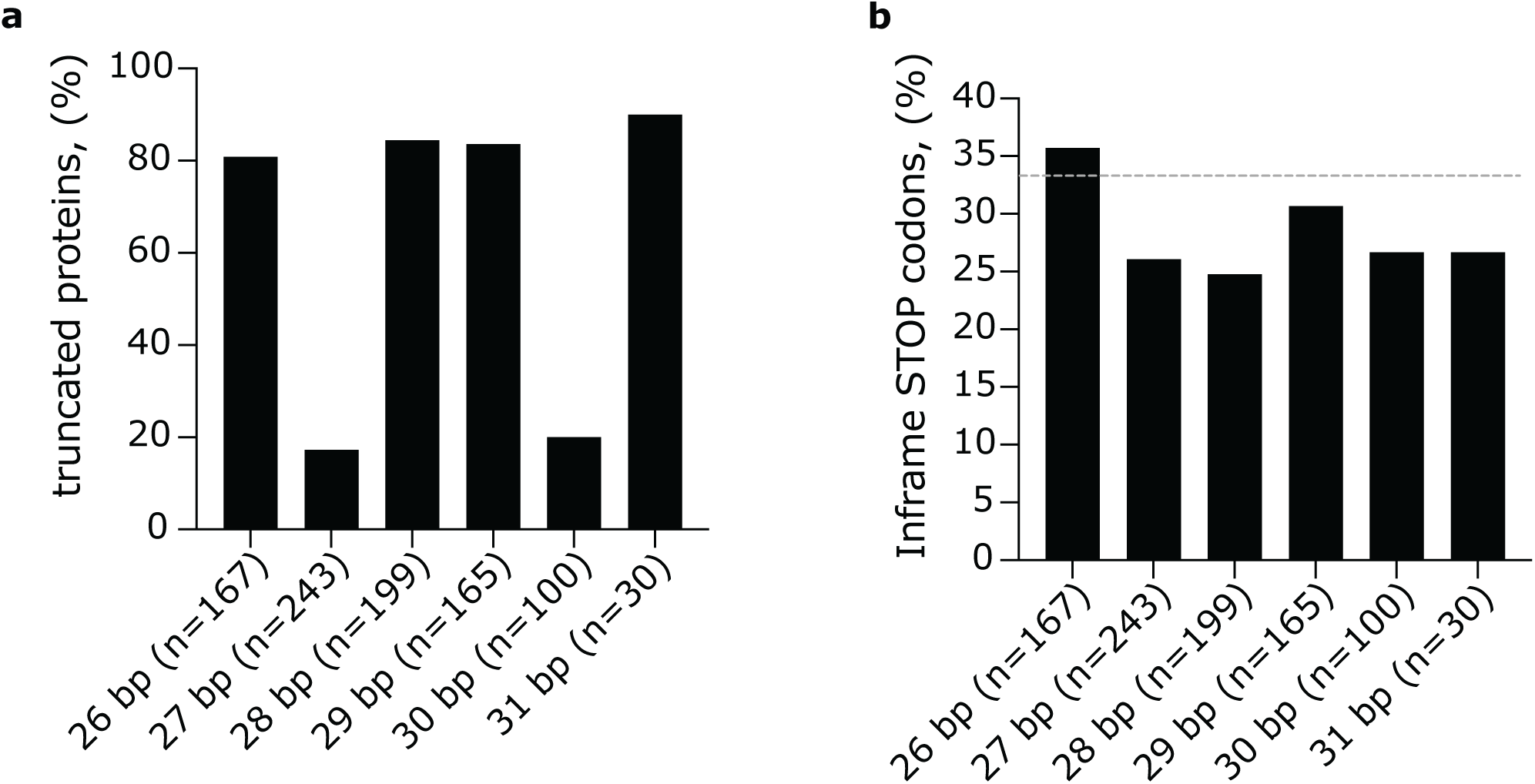
In-frame stop codons are depleted from some of the IES sequences incorporated in transcripts. **a** Percentage of truncated proteins resulting from each insertion sequence length. **b** Fraction of all stop codons in the IES sequences that are in frame and result in a protein truncation. Dashed line shows the expected random rate.

**Supplementary Figure 9.**
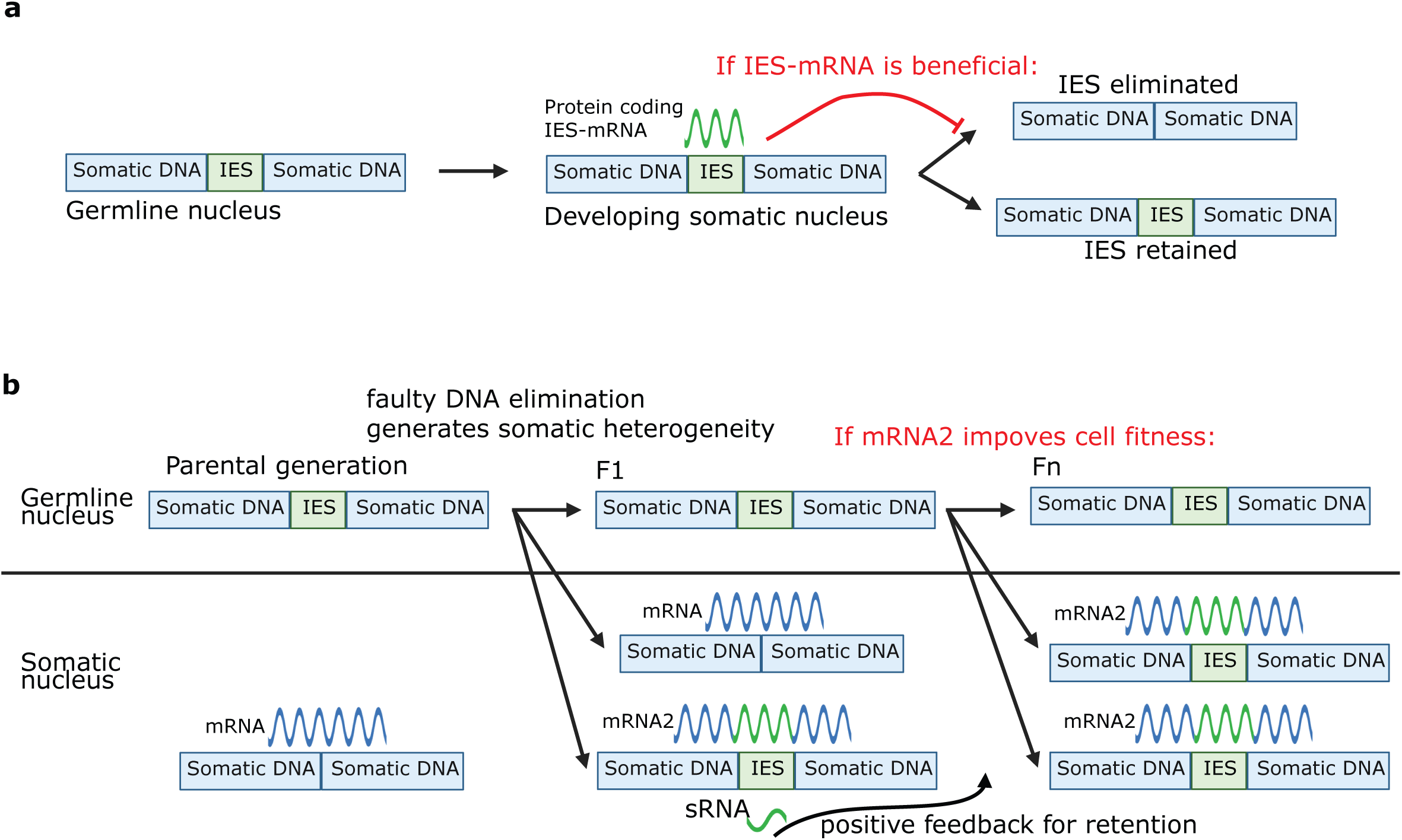
Models for the role of nuclear dualism as enhancer of genome adaptability. **a** A model for IES mediated genome evolution by modified mRNAs expressed during somatic genome development. **b** Generalized model for IES-mediated genome evolution. The F1 generation shows heterogeneity in the somatic nucleus, which can enhance the adaptability of the cell population to various conditions. The somatic nucleus divides amitotically, leading to uneven distribution of the chromosomal variants during vegetative growth.

